# Activation or inhibition of PPARα-mediated fatty acid β-oxidation does not active cardiomyocyte proliferation in normal or infarcted adult mice

**DOI:** 10.1101/667964

**Authors:** Rajika Roy, Tani Leigh, Erhe Gao, Xiaoying Zhang, Ying Tian

**Affiliations:** Department of Pharmacology, Center for Translational Medicine, Lewis Katz School of Medicine, Temple University, Philadelphia, PA 19140, USA; Department of Physiology, Cardiovascular Research Center, Temple University Lewis Katz School of Medicine, Philadelphia, PA 19140, USA

**Keywords:** fatty acid oxidation, cardiomyocyte, proliferation, adult mice

## Abstract

**Objectives:** PPAR genes are known as the important regulators of fatty acid oxidation and energy homeostasis. PPARα is highly expressed in the embryonic and adult heart. Previous studies from infant mouse hearts have suggested that activation of PPARα using GW7647 treatment or cardiac-restricted activation of PPARα using αMHC-PPARα transgenic mice enhanced fatty acid β-oxidation and promoted cardiomyocyte proliferation rate in the postnatal day 4 mouse heart. Here, we further investigate the impact of PPARα-mediated fatty acid β-oxidation on cardiomyocyte proliferation in the adult mouse heart.

**Methods and Results:** Adult wild-type (C57BL/6J) mice were subjected to five injections of GW7647, a highly specific PPARα agonist, or vehicle (saline). Cardiomyocyte proliferation was analyzed by quantification of DNA synthesis via ethynyldeoxyuridine (EdU) incorporation and quantification of cells undergoing mitosis using phosphorylated histone H3 (PH3). GW7647 treatment resulted in activation of PPARα target genes associated with fatty acid metabolism and β-oxidation, validating its biological activity. However, GW7647 treatment did not active cardiomyocyte proliferation in the normal heart. In parallel, mice were subjected to myocardial infarction (MI) using permanent coronary artery occlusion. Both GW7647-treatd wild-type mice and αMHC-PPARα transgenic mice showed no significant differences in cardiomyocyte DNA synthesis and mitosis compared with vehicle-treated wild-type mice after MI. Furthermore, inhibition of PPARα-mediated fatty acid β-oxidation using etomoxir (ETO) treatment had no impact on cardiomyocyte proliferation in both normal and infarcted hearts of wild-type mice compared with vehicle treatment.

**Summary:** These findings suggest that activation or inhibition of PPARα-mediated fatty acid β-oxidation did not active cardiomyocyte proliferation in normal or infarcted hearts of adult mice. Any effects on cardiac function observed following PPARα activation treatment is independent of enhanced cardiomyocyte renewal in the adult heart.

## INTRODUCTION

Heart disease is the number one cause of morbidity and mortality in the Western world. Ischemic injury to cardiac muscle due to acute coronary blockage and other causes can result in a significant loss of cardiomyocytes and decreased cardiac output. This in turn can lead to either death or progressive heart failure. Several approaches for cardiac repair and regeneration have been attempted including replenishment of lost cardiomyocytes through reactivation of pre-existing cardiomyocyte proliferation. Recent studies have discovered the limited potential for regenerative growth in the adult mammalian heart [1–3]. These data have intensified efforts toward stimulating the regeneration of the adult heart muscle from endogenous cardiomyocyte to ameliorate declining cardiac function associated with cardiomyocyte loss and progressive end-stage disease.

The peroxisome proliferator-activated receptors (PPARs) are members of nuclear receptor superfamily [4–6]. There are three PPAR genes, PPARα, PPARβ/δ and PPARγ, each of which is known as the powerful regulator of fatty acid oxidation and energy homeostasis [5, 7, 8]. PPARα is highly expressed in the embryonic and adult heart [7]. Although mice with ablation of PPARα were normal at birth, they developed progressive cardiac fibrosis and reduced cardiac contractile performance in adult life [9, 10]. Furthermore, downregulation of PPARα gene expression was observed in the pressure overload-induced hypertrophied mouse heart [11]. Similarly, decreased PPARα-mediated regulatory pathway activity was observed in the failing human heart [12, 13]. Collectively, these studies indicate that decreased PPARα activity is associated with adverse cardiac function in adult hearts. This view is supported by the observation that increasing PPARα gene expression has a positive impact on cardiomyocytes. For example, treatment with GW7647, a highly specific PPARα agonist that increases fatty acid uptake and β-oxidation [14], increased expression of genes associated with enhanced fatty acid β-oxidation, cardiomyocyte survival and function *in vitro* and *in vivo* [15, 16]. GW7647 treatment led to reduced myocardial infarct size and improved post-ischemic contractile recovery in the *in vivo* and *ex vivo* animal models of ischemia/reperfusion [15, 16]. These findings prompted several clinical trials using PPARα agonists to treat cardiovascular diseases [17–20].

Our previous studies from infant mice have indicated that GW7647 treatment or cardiac-restricted activation of PPARα enhanced fatty acid β-oxidation and promoted cardiomyocyte proliferation rate in the postnatal day 4 mouse heart [21]. Interestingly, recent studies of PPARβ/δ, the closely related family member to PPARα, have suggested that enhanced PPARδ activation promoted cardiomyocyte proliferation and cardiac repair after injury in adult mice [22]. It remains largely unknown what role PPARα plays in cardiomyocyte proliferation and repair from injury in the adult heart.

In this study, we further examined the impact of PPARα activation on cardiomyocyte proliferation during both homeostasis and cardiac repair following myocardial infarction-induced injury in the adult mouse heart. We monitored DNA synthesis in cardiomyocytes using EdU incorporation and quantified cardiomyocytes undergoing mitosis using mitotic marker, phosphor-histone H3 (PH3). Our study showed that PPARα activation using GW7647 treatment increased fatty acid β-oxidation in adult mouse hearts. However, no significant differences in cardiomyocyte proliferation were observed in the heart from GW7647-treated wild-type mice or from αMHC-PPARα transgenic mice compared with the vehicle-treated wild-type mice. In addition, inhibition of PPARα-mediated fatty acid β-oxidation using etomoxir (ETO) treatment had no impact on cardiomyocytes mitosis of infarcted hearts, although there was an increase in cardiomyocytes DNA synthesis compared with vehicle treatment. Furthermore, ETO-induced inhibition of fatty acid β-oxidation resulted in cardiomyocyte dedifferentiation and severe cardiac dysfunction in the infarcted heart compared with vehicle or GW7647 treatment. These results suggest that activation or inhibition of PPARα-mediated fatty acid β-oxidation in adult mouse heart did not active cardiomyocyte proliferation. Any effects on cardiac function observed following PPARα activation treatment is independent of enhanced cardiomyocyte renewal in the adult heart.

## METHODS

### Animal studies

C57BL/6J mice were purchased from the Jackson Laboratory. Generation and genotyping of the αMHC-PPARα line has been previously described [23]. In general, sample size was chosen to use the least number of animals to achieve statistical significance and no statistical methods were used to predetermine sample size. Animals were allocated to experimental groups based on genotype and we did not use exclusion, randomization or blinding approaches. All the animal experiments were performed according to the NIH guidelines (Guide for the care and use of laboratory animals). All experimental procedures involving animals in this study were reviewed and approved by Temple University Medical Center’s Institutional Animal Care and Use Committee.

### Myocardial infarction model

Adult mice (7-8 weeks old, 22-25 gram, male) of the C57BL/6J background were used. Surgically induced myocardial infarction (MI) was performed as previously described [24]. Briefly, after an adequate depth of anesthesia was attained by intraperitoneal injection of Avertin (200-300 mg/kg, i.p,) and/or inhalation of isoflurane (1-3%), the regions of surgical areas were shaved and antiseptic agents (betadine and 70% ethanol) were then applied. A skin incision of 0.8-1.2cm is made along the left side lower margin of pectoral major muscle. After the dissection of pectoral major and minor, the heart will be externalized through 4th intercostal space, and MI will be induced via the ligation of left main descending coronary artery. Heart will then be placed back to the chest and followed by closing of the incision with absorbable sutures. Sutures will be monofilament. Sham operation will be the same as MI procedure except coronary artery will not be occluded. One dose of buprenorphine (0.05-0.1mg/kg) will be given immediately after the procedures. Seven days post-MI, mice were randomly assigned to one of the following groups: MI + vehicle (saline); MI + GW7647; MI + ETO. GW7647 (2 μg/g/day) and ETO (15 μg/g/day) were administrated 1 dose per day for 5 doses by intraperitoneal (i.p.) injection.

### Echocardiography

Mice were anesthetized with inhalation of isoflurane induction 3%, followed by maintenance at 2% using a nose cone. The mouse was placed on a warm platform in the supine position to keep the body temperature around 37°C. The chest hair is removed using hair removal gel cream (Nair). The limbs are taped onto the metal EKG leads. Echo was performed using VisualSonic Vevo 2100 system with a 40 MHz transducer for cardiac imaging. In brief, by placing the transducer along the long-axis of LV, and directing to the right side of the neck of the mouse, two-dimensional LV long-axis is obtained. Then the transducer is rotated clockwise by 90 degree, and the LV short-axis view is visualized. 2D-guided LV M-mode at the papillary muscle level is recorded from either the short-axis view and/or the long-axis view. Trans-mitral inflow Doppler spectra are recorded in an apical 4-chamber view by placing the sample volume at the tip of the mitral valves. Echo images are downloaded and analyzed offline using images analyzing software (Vevo 2100, 1.70, VisualSonic). At least three beats of imaging were measured and averaged for the interpretation of any given measurement. End-diastolic and end-systolic left ventricular internal diameters (LVIDd, LVIDs) were measured from the left ventricular short axis view with 2D orientated M-mode imaging. Left ventricular systolic function was estimated by fractional shortening (FS, %) according to the following formula: FS (%) = [(LVIDd − LVIDs) / LVIDd] × 100. Ejection fraction (EF) was calculated using the end-systolic and end-diastolic volumes as described [25].

### In vivo treatment and EdU labeling

Mice were treated with GW7647 (2 μg/g/day; Sigma, G6793) or etomoxir (15 μg/g/day; Sigma, E1905) or vehicle (saline) via intraperitoneal (i.p.) injection, one dose per day. For EdU labeling, mice were injected with EdU 50 mg/kg via intraperitoneal injection, one dose per day for two doses, and sacrificed at 18 hours after last dose of EdU treatment.

### Histology

Adult heart tissues were fixed in 4% formaldehyde individually and processed for paraffin histology and sectioned using routine procedures. Immunofluorescent staining was performed using previously described protocol [26]. Primary antibodies are: Phospho-Histone H3 (PH3, 1:200; Cell Signaling Technology; 9706L), cardiac Troponin T (cTnT, 1:100; Thermo Scientific, MS-295-P1), Wheat Germ Agglutinin (WGA) Alexa Fluor® 633 Conjugate was used on the same sections to outline cardiomyocytes. DAPI was used to counterstain nuclei. DNA synthesis was measured using Click-iT® EdU (5-ethynyl-2’-deoxyuridine) Alexa Fluor® Imaging Kit (Thermo). The slides were imaged and subjected to an independent blinded analysis, using a Zeiss LSM 710 confocal microscope and ImageJ software. Images shown are representative view of multiple fields from at least four independent samples per group. Quantitation of cell numbers was done using images acquired on confocal microscopy and the ImageJ with the “Cell Counter” plug-in, counting multiple fields from at least 4 independent samples per group and about 3200 −7837 cTnT+ cells per sample.

### RNA purification and qRT-PCR analysis

Quantitative real-time PCR (qRT-PCR) analysis was performed using Trizol isolated RNA, which was used to generate cDNA using random hexamer primers and SuperScript III RT (Invitrogen). qRT-PCR primer sequences are listed in Table 1. SYBR green detection of amplification was performed using the StepOne Plus cycler (Applied Biosystems). Transcript expression values were generated with the comparative threshold cycle (Delta CT) method by normalizing to the expression of the 18S gene.

**Table 1.**
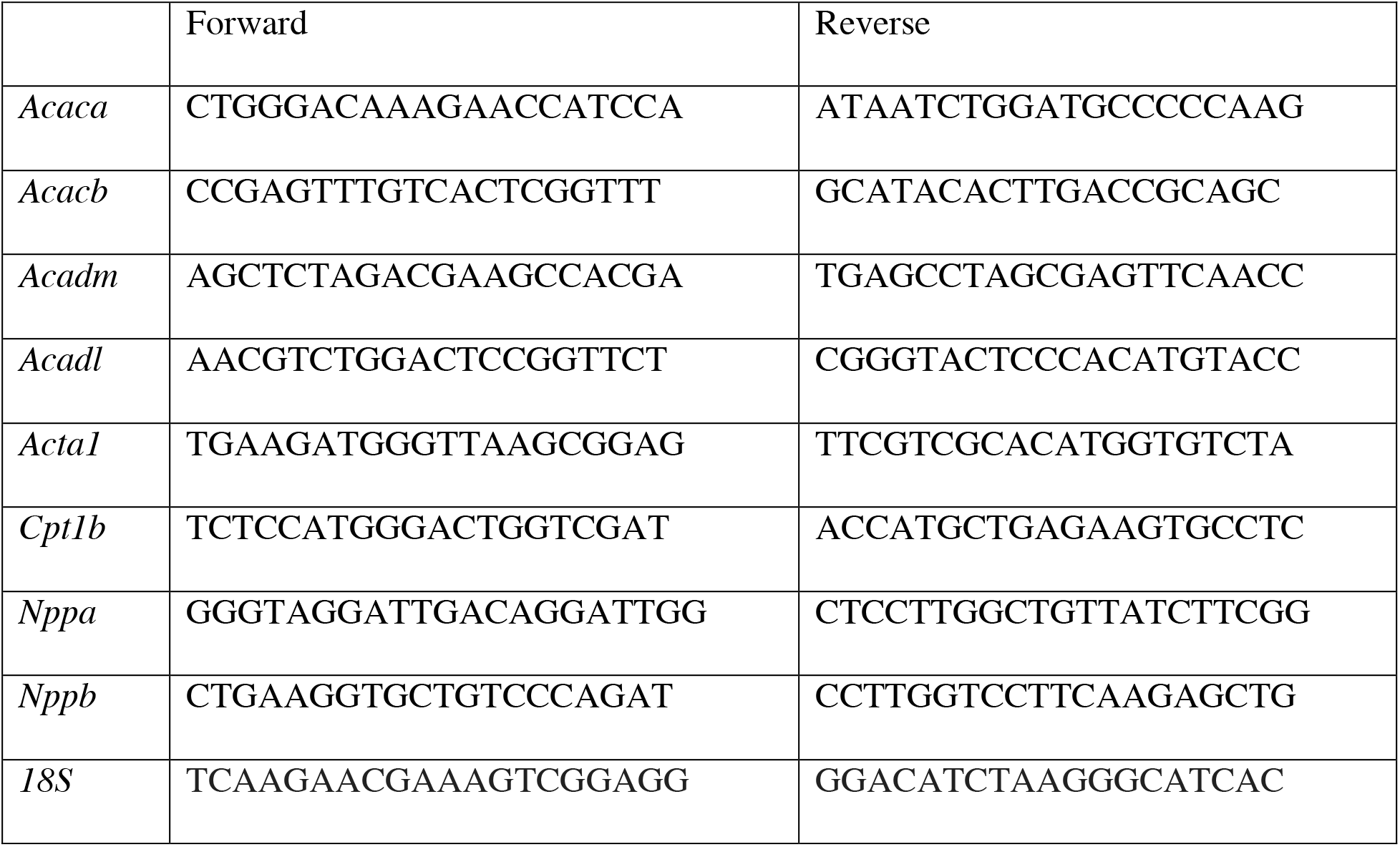
qRT-PCR primer sequences used in this study.

### Statistical analysis

Data are presented as mean ± standard error of the mean (s.e.m.). Student’s t-test and one-way ANOVA were used to calculate statistical significance. *P* values are depicted as follows: * *P* < 0.05; ** *P* < 0.01; *** *P* < 0.001; **** *P* < 0.0001. Results with *P* > 0.05 were considered not significant (*n.s*.). All analyses were performed with GraphPad Prism 7.

## RESULTS

C57BL/6J mice were administered with GW7647 for a total of 5 doses by i.p injections (Figure 1A). The control mice received vehicle (saline) only. Heart ventricles were harvested 24 hours after the last injection. Previous studies demonstrated the expression of fatty acid metabolism-associated genes (*Acaca*, *Acacb*, *Acadl*, *Acadm*, *Cpt1b*) as indicators of PPARα biological activity *in vivo* [21]. Consistent with these previous studies, GW7647 treatment resulted in a significant increase in the mRNA level of these genes (Figure 1B). EdU incorporation was used to monitor cardiomyocyte DNA synthesis. At 42 and 18 hours before harvesting the heart, EdU was injected 6 hours after GW7647 treatment (Figure 1A). Cardiomyocyte DNA synthesis was identified by using Click-iT EdU Alexa Fluor and co-immunostaining of anti-cardiac troponin T in heart ventricular tissue sections. No significant difference in cardiomyocyte DNA synthesis was detected in GW7647-treated mice compared with saline-treated mice (Figure 1C). To determine if inhibition of PPARα-mediated activation of cardiac fatty acid β-oxidation had impact on cardiomyocyte DNA synthesis, C57BL/6J mice were given etomoxir (ETO), an inhibitor of carnitine palmitoyltransferase I (CPT1) [27], which is a key regulator of mitochondrial fatty acid uptake, in a similar approach as GW7647 treatment (Figure 1A). ETO-treated mice had a significant reduction of mRNA level of fatty acid metabolism-associated genes (*Acaca*, *Acacb*, *Acadl*, *Acadm*, *Cpt1b*) compared to the saline-treated mice (Figure 1B). However, no difference in cardiomyocyte DNA synthesis was observed in mice with ETO treatment compared with saline treatment (Figure 1C). Collectively, these results indicated that GW7647-induced activation or ETO-induced inhibition of PPARα-mediated cardiac fatty acid β-oxidation did not active cardiomyocyte into cell cycle during homeostasis in the adult heart.

**Figure 1.**
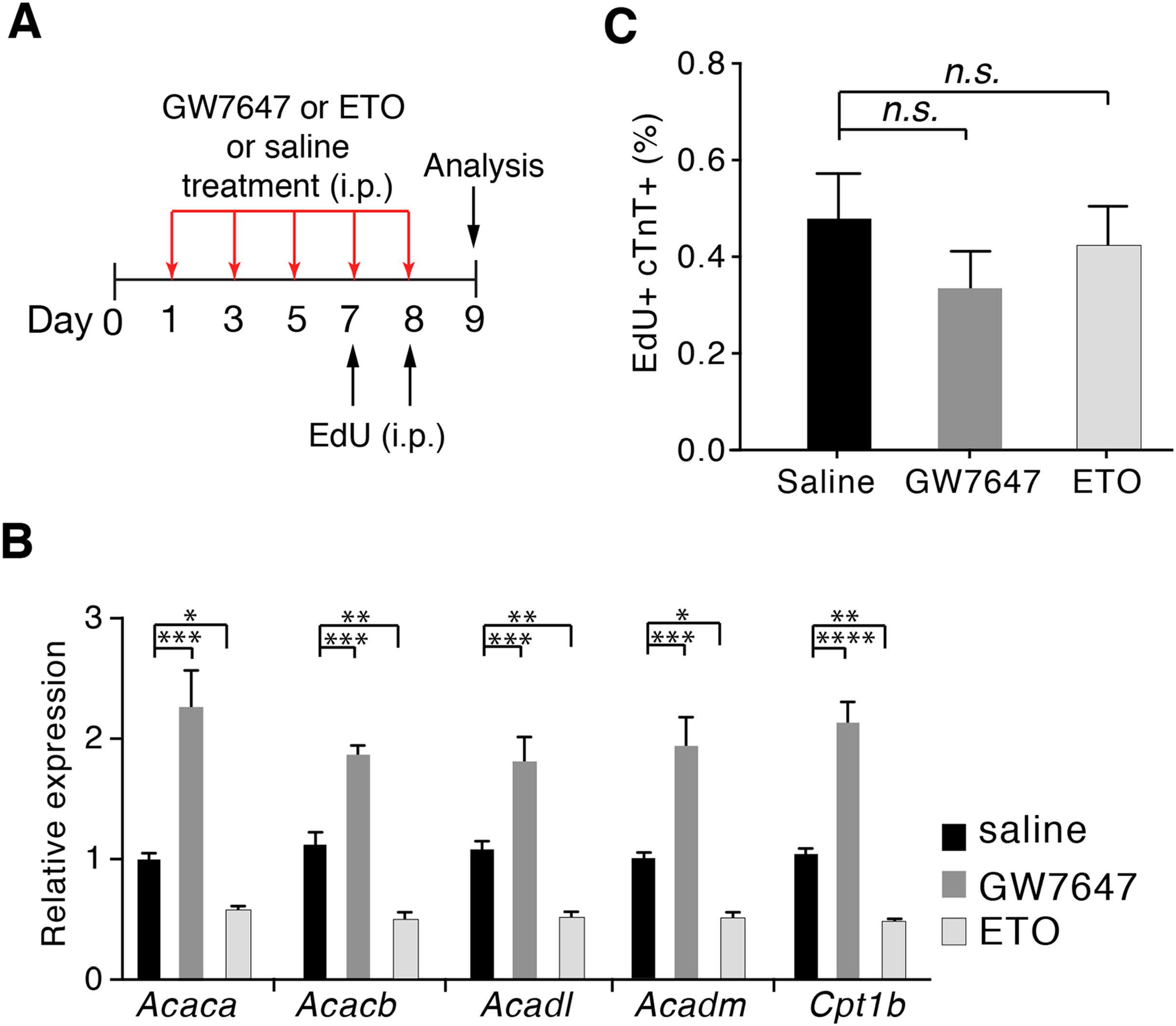
Effects of fatty acid oxidation on cardiomyocyte proliferation in normal adult hearts. (A) Schematic of experimental design. (B) Quantification of PPARα target genes associated with fatty acid metabolism by qRT-PCR analysis of the mRNA of isolated heart ventricles (n=4-5 per group). (C) Quantification of cardiomyocyte DNA synthesis using EdU+cTnT+ cells as percentage of total cTnT+ cells analyzed on heart ventricular tissue sections (n=4 per group). \**P* < 0.05; \*\**P* < 0.01; \*\*\**P* < 0.001; \*\*\*\**P* < 0.0001, one-way ANOVA.

It has been shown that adult cardiomyocytes exhibit increased cell cycle activity in response to myocardial infarction (MI) induced cardiac injury [28–30]. To determine whether GW7647 or ETO treatment had impact on cardiomyocyte proliferation during cardiac repair following injury, C57BL/6J mice were treated with GW7647 or ETO, using i.p. injections for a total of 5 doses during 7-14 days after MI (Figure 2A), the time window when increased cell cycle activity has been observed in cardiomyocytes [28–30]. Heart ventricles were analyzed 24 hours after the last injection. We observed a significant increase in the number of cardiomyocyte incorporating EdU in the left ventricle and septum of the infarcted mice compared with sham mice (Figure 2B), consistent with previous studies [28–30]. However, no significant difference in cardiomyocyte DNA synthesis was detected in mice receiving GW7647 as compared to mice receiving vehicle (saline) alone. In contrast, ETO treatment resulted in a significant increase in cardiomyocyte incorporating EdU (3.8% ± 0.9% vs. 1.5% ± 0.3%, ETO vs. saline treatment, *P*<0.05, Figure 2B). Furthermore, analysis of cardiomyocytes in mitotic phase using mitotic cell cycle marker phosphorylated histone H3 (PH3) showed a significant increase in the number of PH3+ cardiomyocytes (PH3+cTnT+) in the infarcted heart compared with the sham heart (Figure 2C). However, there were no significant differences in cardiomyocytes undergoing mitosis between GW7647-, ETO- and saline-treated mice, although there was a trend towards a reduced PH3+ cardiomyocytes in the ETO-treated mice (Figure 2C).

**Figure 2.**
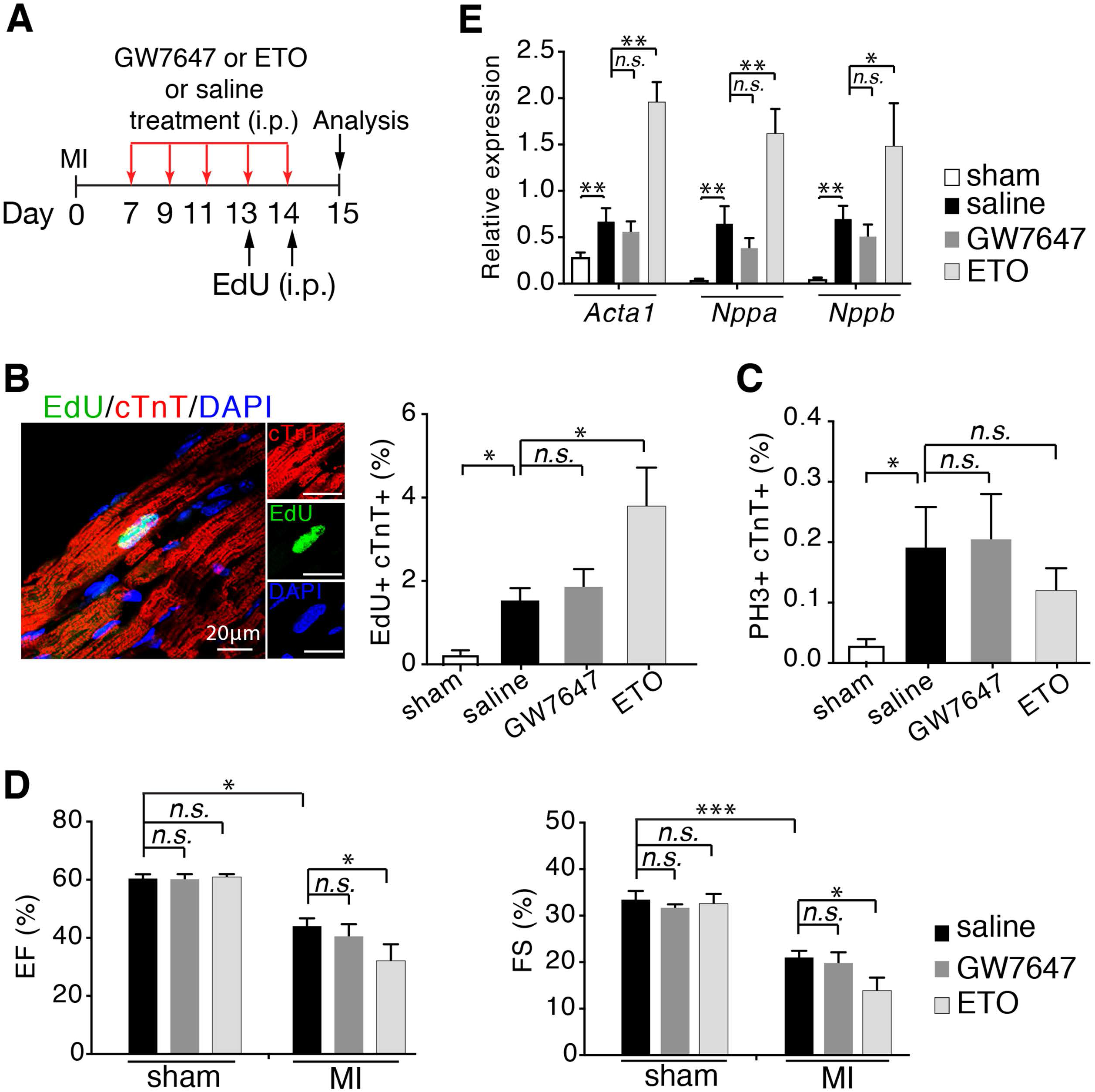
Effects of fatty acid oxidation on cardiomyocyte proliferation in adult hearts following myocardial infarction (MI). (A) Schematic of experimental design for studies performed in (B-E). (B) Cardiomyocytes in DNA synthesis-phase were detected by using Click-iT EdU Alexa Fluor (green) and co-immunostaining with antibody against cTnT (red) on tissue sections. Quantification of EdU+cTnT+ cells as percentage of total cTnT+ cells analyzed per field (n=5 per group). Scale bars: 20μm. (C) Quantification of cardiomyocytes in mitotic phase using PH3+cTnT+ cells as percentage of total cTnT+ cells analyzed on heart ventricular tissue sections (n=5 per group). (D) Cardiac function in mice evaluated by echocardiography at days 15 (D15) post-MI (n=5-13 per group). EF, ejection fraction; FS, fractional shortening. (E) Quantification of genes associated with fetal cardiomyocyte gene program by qRT-PCR analysis of the mRNA of isolated heart ventricles (n=4-5 per group). \**P* < 0.05; \*\**P* < 0.01; \*\*\**P* < 0.001, one-way ANOVA (B-E) and Student’s t test (B, C, E).

Cardiac functional assessment revealed a significant reduction in the levels of ejection fraction (EF) and fractional shortening (FS) of the infarcted mice compared with sham mice (Figure 2D). No differences in these functional assessments were observed in GW7647-treated animals compared with saline-treated animals. However, ETO treatment led to significant lower levels of EF and FS compared with the saline treatment (Figure 2D). Our previous studies have suggested that ETO treatment prevented cardiomyocyte maturation in early postnatal mouse hearts [21]. We surmised that ETO treatment might induce cardiomyocyte dedifferentiation in the adult heart after MI. To test this possibility, we examined the expression of genes associated with cardiomyocyte maturation at 24 hours after the last dose of ETO injection. Analysis of heart ventricles by qRT-PCR showed a significant up-regulation of *Acta1*, *Nppa* and *Nppb*, the markers for fetal gene program in mammalian hearts that are associated with cardiac dysfunction [31], in ETO-treated hearts compared with saline-treated hearts (Figure 2E). In contrast, GW7647 treatment had no impact on the expression of these gene compared with saline treatment (Figure 2E). To determine whether GW7647 or ETO treatment affected cardiac fibrosis formation and heart function over the long term, we examined mouse hearts at 58 days post-MI (Figure 3A). No significant differences in fibrotic scarring were observed between GW7647-, ETO-, and saline-treated mice (Figures 3B-C). Cardiac functional assessment showed animals with GW7647 treatment had similar levels of EF and FS in infarcted hearts compared with saline treatment (Figures 3D-F). In contrast, ETO-treated animals exhibited severe cardiac dysfunction as evidenced by the significant decreases in EF and FS in infarcted hearts compared with saline-treated animals (Figures 3D-F).

**Figure 3.**
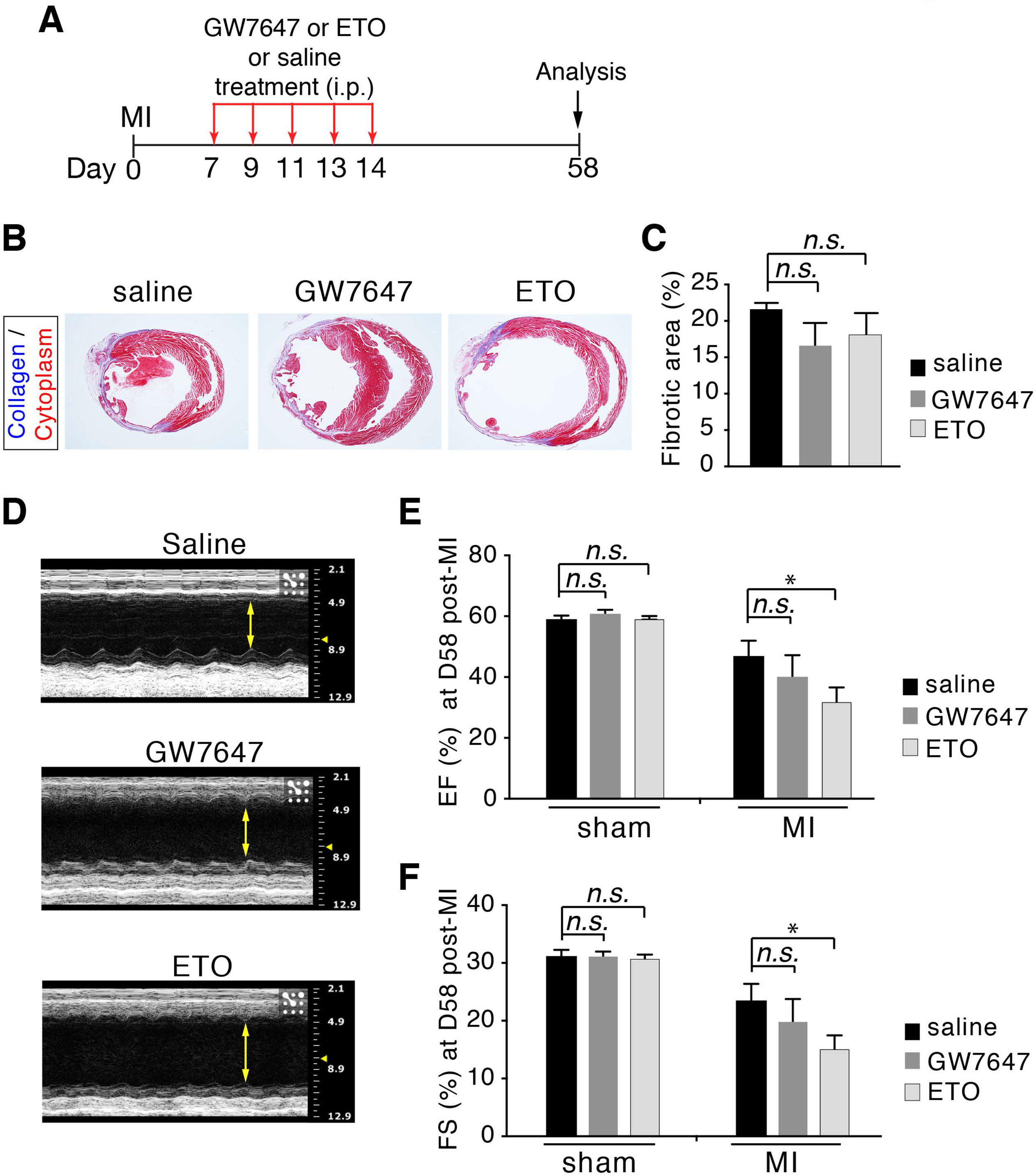
Effects of fatty acid oxidation on cardiac fibrosis and function following myocardial infarction (MI). (A) Schematic of experimental design for studies performed in (B-F). (B) Masson’s trichrome-stained heart sections at days 58 post-MI. (C) Quantification of fibrotic regions in heart sections (n=5 per group). (D-F) Cardiac function in mice evaluated by echocardiography at days 58 post-MI (n=5 per group). EF, ejection fraction; FS, fractional shortening. \**P* < 0.05, one-way ANOVA.

To further confirm PPARα-mediated increase in fatty acid β-oxidation in cardiomyocytes, we utilized a previously established transgenic mouse line, in which PPARα is overexpressed specifically in cardiomyocytes (αMHC-PPARα) [32]. Adult αMHC-PPARα mice (7-week-old) were subjected to MI. Thirteen days later, mice received two daily EdU injections and hearts were harvested 24 hours later and processed (Figure 4A). Consistent with the findings observed with the GW7647 treatment, no significant difference in cardiomyocytes DNA synthesis (EdU+cTnT+) was observed between αMHC-PPARα mice and the wild-type littermates (Figure 4B). At 30 days after MI, αMHC-PPARα mice showed similar levels of cardiac fibrosis formation compared with wild-type littermates (Figures 4C-D). Moreover, no significant differences in cardiac functions (EF and FS) were detected between αMHC-PPARα mice and the wild-type littermates (Figures 4E-F). Altogether, these results indicated that activation of PPARα-mediated fatty acid β-oxidation did not active cardiomyocyte proliferation in both normal and infarcted hearts of adult mice. Inhibition of fatty acid β-oxidation using ETO had no impact on cardiomyocyte proliferation in the adult mouse. ETO treatment led to cardiomyocyte dedifferentiation and severe cardiac dysfunction in the infarcted heart.

**Figure 4.**
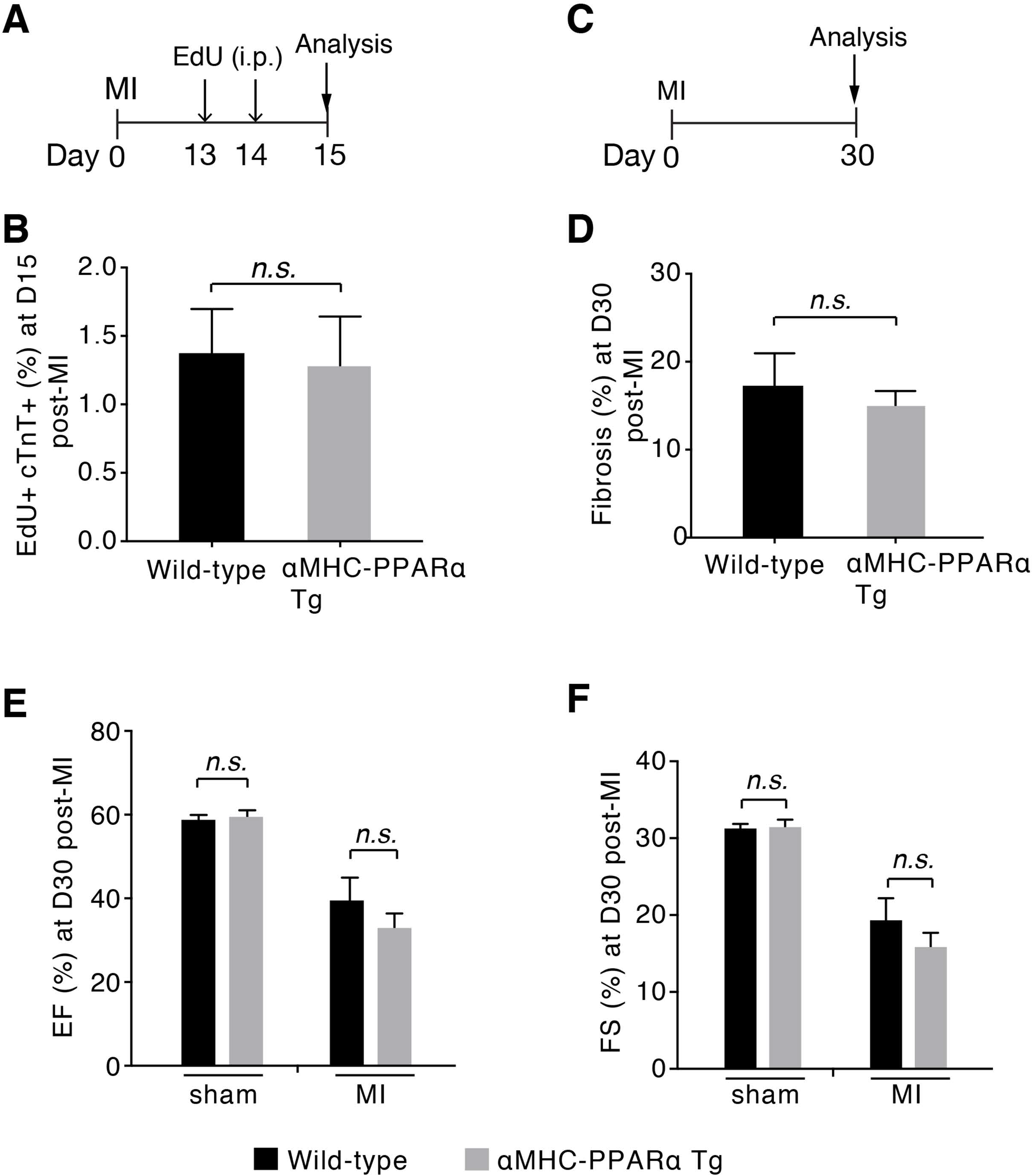
Analysis of cardiomyocyte proliferation, cardiac fibrosis and function in αMHC-PPARα transgenic (Tg) mice after myocardial infarction (MI). (A) Schematic of experimental design for studies performed in (B). (B) Quantification of EdU+cTnT+ cells as percentage of total cTnT+ cells analyzed on heart ventricular tissue sections (~5000 cTnT+ cells per sample) at days 15 post-MI (n=4-5 per group). (C) Schematic of experimental design for studies performed in (D-F). (D) Quantification of fibrotic regions in heart sections at days 30 post-MI (n=8-10 per group). (E and F) Cardiac function in mice evaluated by echocardiography at days 30 post-MI (n=6-8 per group). EF, ejection fraction; FS, fractional shortening. Student’s t test.

## DISCUSSION

Previous studies have demonstrated that the mouse and human heart exhibits a very low level of adult cardiomyocyte proliferation, which can be increased after myocardial injury [32, 33]. Bioenergy balance is a key aspect of the perfect operation of various biological systems and cellular processes. In this study, we have assessed the role of cardiac energy metabolism in regulating cardiomyocyte proliferation during adult life. Cell proliferation, differentiation and metabolism are three fundamental features of mammals. Interestingly, cell proliferation and metabolism have both been implicated in other biological processes, such as regulation of pluripotent stem cell state and T cell activation in immune system [34–38]. The studies reported here showed that activation or inhibition of PPARα-mediated fatty acid β-oxidation in the adult mouse heart did not active cardiomyocyte proliferation in both normal and infarcted hearts.

It’s now well established that the fetal and neonatal mammalian heart consumes primarily glucose, and fatty acids become the major fuel for ATP synthesis in cardiomyocytes of adult hearts [39]. Cardiomyocytes proliferate rapidly during fetal life but lose their ability to proliferation soon after birth [40]. However, before terminal withdrawal from the cell cycle, cardiomyocytes undergo another round of cell cycle during early postnatal life in mice, and the neonatal heart can regenerate through increased cardiomyocyte proliferation [41]. This ability to regeneration in response to injury ends by 7 days after birth in mice, corresponding to the exit of cardiomyocyte from the cell cycle. Notably, those changes coincide with the metabolic shift toward fatty acid β-oxidation [21]. Whether cardiac metabolism contributes to cardiomyocyte proliferation in physiological and pathological settings remains unanswered. Our previous studies have suggested that activation of PPARα-mediated fatty acid β-oxidation promoted cardiomyocyte proliferation rate in infant mouse hearts during early postnatal life [21]. However, it is unclear whether PPARα activation has impact on adult cardiomyocyte proliferation. A recent report has suggested that PPARδ, the closely family member of PPARα, regulates cardiomyocyte proliferation and activation of PPARδ promotes cardiac repair following myocardial injury [22]. The studies reported here indicated that activation of PPARα-mediated fatty acid β-oxidation in the adult mouse heart did not active cardiomyocytes proliferation during both homeostasis and cardiac repair following MI-induced injury. Thus, our results in PPARα indicated a distinct concept and suggested PPARα-mediated increase in fatty acid β-oxidation had no impact on cardiomyocyte proliferation in the adult mouse heart. Additionally, earlier studies have demonstrated that activation of PPARα in adult mice protects the heart from ischemia/reperfusion injury in part due to PPARα-mediated increase in fatty acid β-oxidation [15, 16]. Our results suggested that any effects on cardiac function observed following activation of PPARα-mediated fatty acid β-oxidation take place independently of enhanced cardiomyocyte proliferation in the adult heart.

Decreased PPARα-mediated regulatory pathway activity has been observed in the failing human heart [12,13]. Studies reported here showed that inhibiting fatty acid β-oxidation using ETO treatment did elicit some adverse long-term impacts as evidenced by cardiomyocyte dedifferentiation and reduced cardiac function in infarcted mouse hearts compared with either vehicle or GW7647 treatment.

## CONCLUSIONS

In summary, the current findings indicate that activation or inhibition of PPARα-mediated fatty acid β-oxidation did not active cardiomyocyte proliferation in both normal and infarcted hearts of adult mice. Inhibition of fatty acid β-oxidation using ETO treatment caused cardiomyocyte dedifferentiation and severe cardiac dysfunction in the infarcted heart of adult mice. These results suggest that any effects on cardiac function observed following PPARα activation treatment occur independently of enhanced cardiomyocyte renewal in the adult heart.

## ACKNOWLEDGEMENTS

We thank Dr. Brian N. Finck for sharing αMHC-PPARα mice.

## Data Availability

All data generated and analyzed during this study are included in this published article.

## AUTHOR CONTRIBUTIONS

R.R. performed echocardiography in adult mice after cardiac surgery and analyzed data. T.L. performed cell counting and quantification in adult mice. E.G. performed cardiac surgery. X.Z. performed echocardiography in adult normal mice and analyzed data. Y. T. designed the study and wrote the manuscript.

